# Quorum sensing orchestrates parallel cell death pathways in *Vibrio cholerae* via Type 6 secretion dependent and independent mechanisms

**DOI:** 10.1101/2024.09.23.614608

**Authors:** Ameya A. Mashruwala, Bonnie L. Bassler

## Abstract

Quorum sensing (QS) is a cell-to-cell communication process that enables bacteria to coordinate group behaviors. In *Vibrio cholerae* colonies, a program of spatial-temporal cell death is among the QS-controlled traits. Cell death occurs in two phases, first along the colony rim, and subsequently, at the colony center. Both cell death phases are driven by the type VI secretion system (T6SS). Here, we show that HapR, the master QS regulator, does not control *t6ss* gene expression nor T6SS-mediated killing activity. Nonetheless, a Δ*hapR* strain displays no cell death at the colony rim. RNA-Seq analyses reveal that HapR activates expression of an operon containing four genes of unknown function, *vca0646-0649*. Epistasis and overexpression studies show that two of the genes, *vca0646* and *vca0647*, are required to drive cell death in both a Δ*hapR* and a Δ*hapR* Δ*t6ss* strain. Thus, *vca0646*-*0649* are regulated by HapR but act independently of the T6SS machinery to cause cell death, suggesting that a second, parallel pathway to cell death exists in *V. cholerae*.

**Significance:** Cell death is a fundamental biological process. In mammals, cell death sculpts tissues during development, enables injury recovery, and regulates immunity. In bacteria, cell death mechanisms remain little explored. Recently, colonies formed by the pathogen *Vibrio cholerae* were demonstrated to undergo a spatio-temporal program of cell death. The program is controlled by quorum sensing (QS) and driven by the Type VI secretion system. Here, we discover QS-controlled genes, called *vca0646-0649*, that cause cell death in *V. cholerae* colonies independently of the Type VI secretion system. These findings indicate that a second cell death pathway exists in *V. cholerae*. The results expand our understanding of bacterial cell death mechanisms and provide insight into how cell death shapes bacterial community structure.

## Introduction

Quorum sensing (QS) is a process of bacterial cell-cell communication that depends on the production, release, accumulation, and detection of extracellular signal molecules called autoinducers (AIs) (1, 2). QS enables bacteria to monitor the vicinal cell density and coordinate population-wide gene expression and collective behaviors (1, 2). In so doing, bacteria accomplish tasks that require many cells acting in synchrony to make the tasks successful. In the model QS bacterium and pathogen *Vibrio cholerae*, which causes the cholera disease, information encoded in AIs is relayed through two QS pathways both of which converge on a shared transcription factor, LuxO (3). At low cell density (LCD), when AIs are absent, LuxO is phosphorylated (LuxO∼P) and it activates transcription of genes encoding four small RNAs, called Qrr1-4 (4, 5). Qrr1-4 repress production of the HapR transcription factor (5). HapR is the master high cell density (HCD) QS regulator. At HCD, when AI concentrations are above the threshold required for detection, LuxO is dephosphorylated, production of Qrr1-4 ceases, HapR is produced, and it activates expression of genes specifying group behaviors.

The bacterial type VI secretion system (T6SS) is a contact-dependent nanomachine that delivers toxic molecules into other cells (6–8). Briefly, T6SS structural proteins assemble into a syringe-like device, the tip of which is loaded with toxic effector proteins (7, 9). The apparatus injects the effectors into neighboring competitor cells, which kills them. To avoid self-harm, T6SS-active bacteria produce immunity proteins that inactivate the toxic effector proteins (10). Protection from incoming T6SS attacks is also conferred by physical means including exopolysaccharide or capsular polysaccharide macromolecules that act as “shields” (11, 12). In *V. cholerae*, the genes encoding T6SS components are arranged in one large and three auxiliary clusters (Figure S1). Regulation of the T6SS machinery is strain specific, and important for this work is that unlike the commonly studied pandemic El Tor strain, the El Tor *V. cholerae* environmental isolate called 2740-0 expresses its *t6ss* genes under laboratory settings due to the presence of an activating, cis-acting, single nucleotide polymorphism (13, 14). In *V. cholerae*, T6SS function is also QS regulated (15). At LCD, *t6ss* expression from the large cluster is repressed by the Qrr sRNAs via a post-transcriptional mechanism. In addition, the Qrr sRNAs indirectly repress expression of auxiliary *t6ss* clusters by preventing HapR production (15). HapR is an activator of auxiliary *t6ss* gene cluster expression. Simultaneous to reducing T6SS offensive capacity, at LCD, the Qrr sRNAs promote increased production of the Vibrio polysaccharide (Vps) “shield” that blocks incoming T6SS attacks, and thus, boost T6SS defenses (16, 17).

Certain bacteria, including *V. cholerae* 2740-80, form colonies that, over time, develop outgrowths called sectors (16, 18–20). In *V. cholerae*, sector formation is preceded by a cell death program that occurs in two phases (16). The first phase of death occurs at the colony rim and the second phase in the colony center. Relevant to the present work is that cell death at the colony rim is a consequence of T6SS-dependent kin-killing (16). Killing imposes a selective pressure for the bacteria to acquire mutations that enable them to resist killing. As a consequence, these “variants” form the outgrowths called sectors. The *V. cholerae* 2740-80 sector variants commonly possess gain-of-function mutations in *luxO* that “lock” the cells into the QS LCD gene expression program (16). The “locked” *luxO* LCD mutations confer growth advantages by two mechanisms: First, they reduce *t6ss* expression and thus suppress overall T6SS-mediated killing activity. Second, they increase Vps production, which enhances defense against incoming T6SS attacks. Isolation and streaking of the *luxO* variants as pure colonies show that they display no cell death at the colony rim, and they do not sector. However, cell death in the center of the colony continues to occur. Thus, killing at the rim must be a HCD QS-controlled T6SS-dependent trait (16).

Some *V. cholerae* 2740-80 variants isolated from sectors have mutations in *hapR* (Supplementary Table 1) (16). With one exception, the *hapR* mutations confer attenuation or loss-of-function and thus, analogous to the above *luxO* mutations, “lock” the cells into the QS LCD mode. Only one variant, encoding HapR A52T, did not fit this pattern (16). HapR A52T is known to drive expression of both HCD and LCD QS genes (16, 21). The role of HapR or the HapR variants in modulating T6SS function and/or the rim cell death program was not analyzed in the previous study (16). Exploring the role HapR plays in driving *V. cholerae* cell death pattern formation is the topic of this work.

Surprisingly, we discover that, in *V. cholerae* 2740-80, HapR does not regulate *t6ss* gene expression nor T6SS-mediated killing activity. Rather, QS control relies only on the LuxO-Qrr arm of the circuit. Despite being proficient in T6SS-mediated killing, a Δ*hapR* strain nonetheless displays an absence of cell death at the colony rim. RNA-Seq demonstrated that expression of the *vca0646*-*vca0649* operon was diminished in the Δ*hapR* strain. Restoration of *vca0646*-*vca0649* operon expression reestablishes cell death at the colony rim. Introduction of each gene and combination of genes from the operon into a Δ*hapR* strain showed that the *vca0646-0647* pair of genes is sufficient to drive the cell death pattern. VCA0647 was previously identified as a potential repressor of T6SS defense function in *V. cholerae* (22). The obvious hypothesis was that in *V. cholerae* 2740-80, HapR activates *vca0646*-*0649* expression and VCA0646 and VCA0647, in turn, suppress T6SS defense function. Together, these regulatory arrangements enable T6SS kin-killing and cell death to occur at the colony rim. However, again to our surprise, expression of *vca0646*-*0649* restored the cell death pattern in a Δ*hapR* Δ*t6ss* strain that lacks all T6SS killing machinery. Thus, VCA0646 and VCA0647 do not carry out their functions via a T6SS-mediated mechanism. While overexpression of *vca0646-0649* promoted cell death, deletion of these genes did not alter the cell death pattern. This finding suggests redundant or additional components exist that can compensate for loss of *vca0646-0649*. We conclude that VCA0646-0647 participate in a new QS-regulated, T6SS-independent cell death pathway in *V. cholerae* (Figure 1).

**Figure 1.**
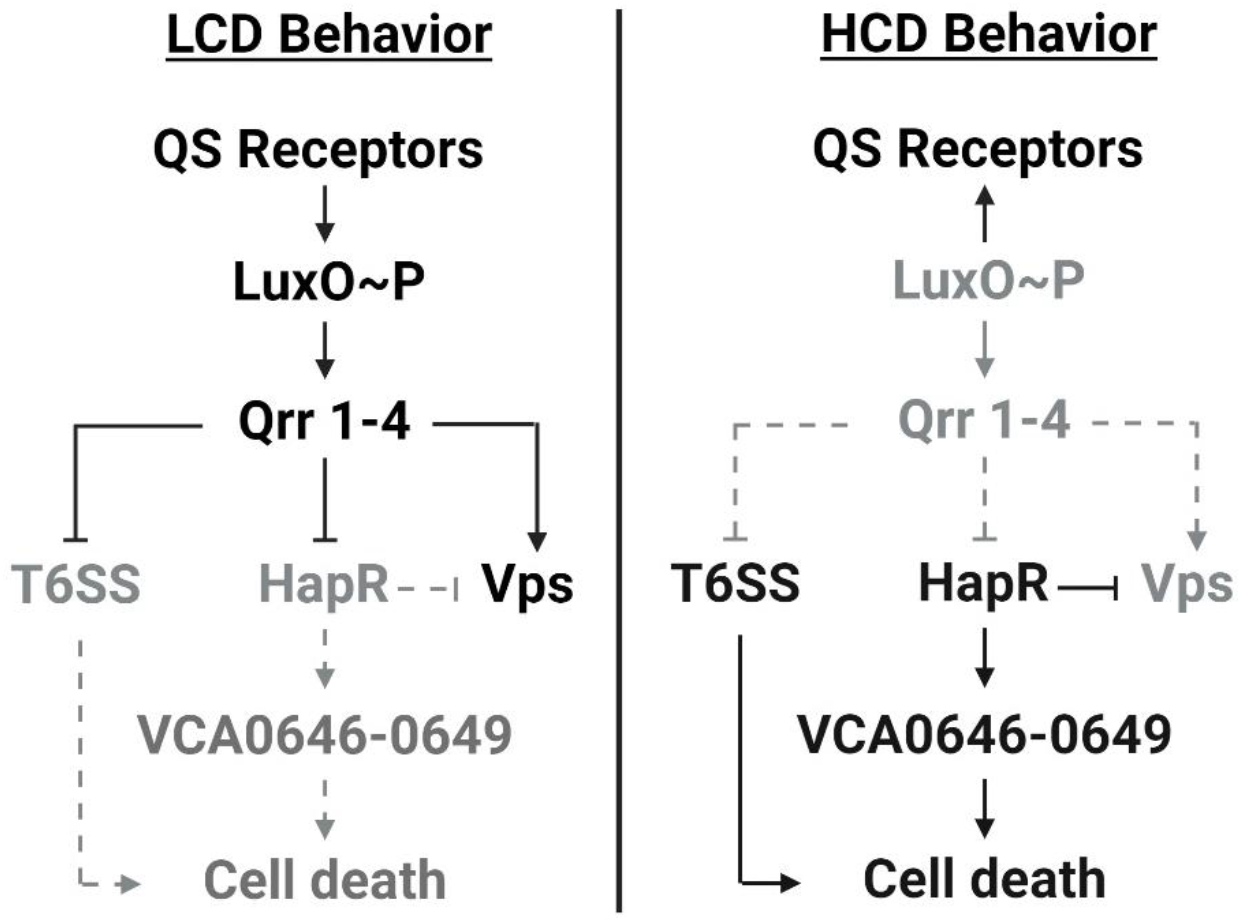
Simplified model of *V. cholerae* 2740-80 QS regulation of *t6ss, vps*, and *vca0646-0649*, and consequently, the HCD-specific cell death behavior.

## Results

### QS control of T6SS-mediated killing activity in *V. cholerae* 2740-80 is driven by LuxO∼P and the Qrr sRNAs independently of HapR

It was previously reported that some variants recovered from *V. cholerae* 2740-80 colony sectors had acquired mutations in *hapR* (Supplementary Table 1) (16). However, the mechanism connecting the *hapR* mutations to T6SS-mediated cell death was not investigated. We do that here starting by assessing whether *V. cholerae* 2740-80 Δ*hapR* or *V. cholerae* 2740-80 possessing the *hapR* variant mutations display altered *t6ss* gene expression compared to wildtype (WT) *V. cholerae* 2740-80. To measure expression, we constructed a luciferase (*lux*) transcriptional fusion to the *hcp2* promoter (designated *hcp2*-*lux*). Hcp2 is encoded by the first gene in the T6SS operon that also harbors *vasX* (Figure S1). VasX is a key T6SS toxin that drives cell death at the rim of *V. cholerae* 2740-80 colonies (16, 22). To avoid complications from possible secondary mutations in the *hapR* variants originally obtained from colony sectors, we reintroduced each *hapR* allele from the variants into the parental *V. cholerae* 2740-80 strain. As a control, we included *V. cholerae* 2740-80 carrying *luxO A97E* in our analyses (16). LuxO A97E is a phosphomimetic allele that confers the QS LCD state (16). *V. cholerae* 2740-80 *luxO A97E* has decreased expression of *t6ss* genes (16). Indeed, when strains were grown to HCD, *V. cholerae* 2740-80 *luxO A97E* displayed ∼15-fold lower *hcp2*-*lux* activity than WT *V. cholerae* 2740-80 (Figure 2A). By contrast, at HCD, the Δ*hapR* strain and the strains harboring the variant *hapR* alleles did not exhibit altered *hcp2*-*lux* expression, producing light levels similar to that of *V. cholerae* 2740-80 at HCD (Figure 2A and Supplementary Table 1).

**Figure 2.**
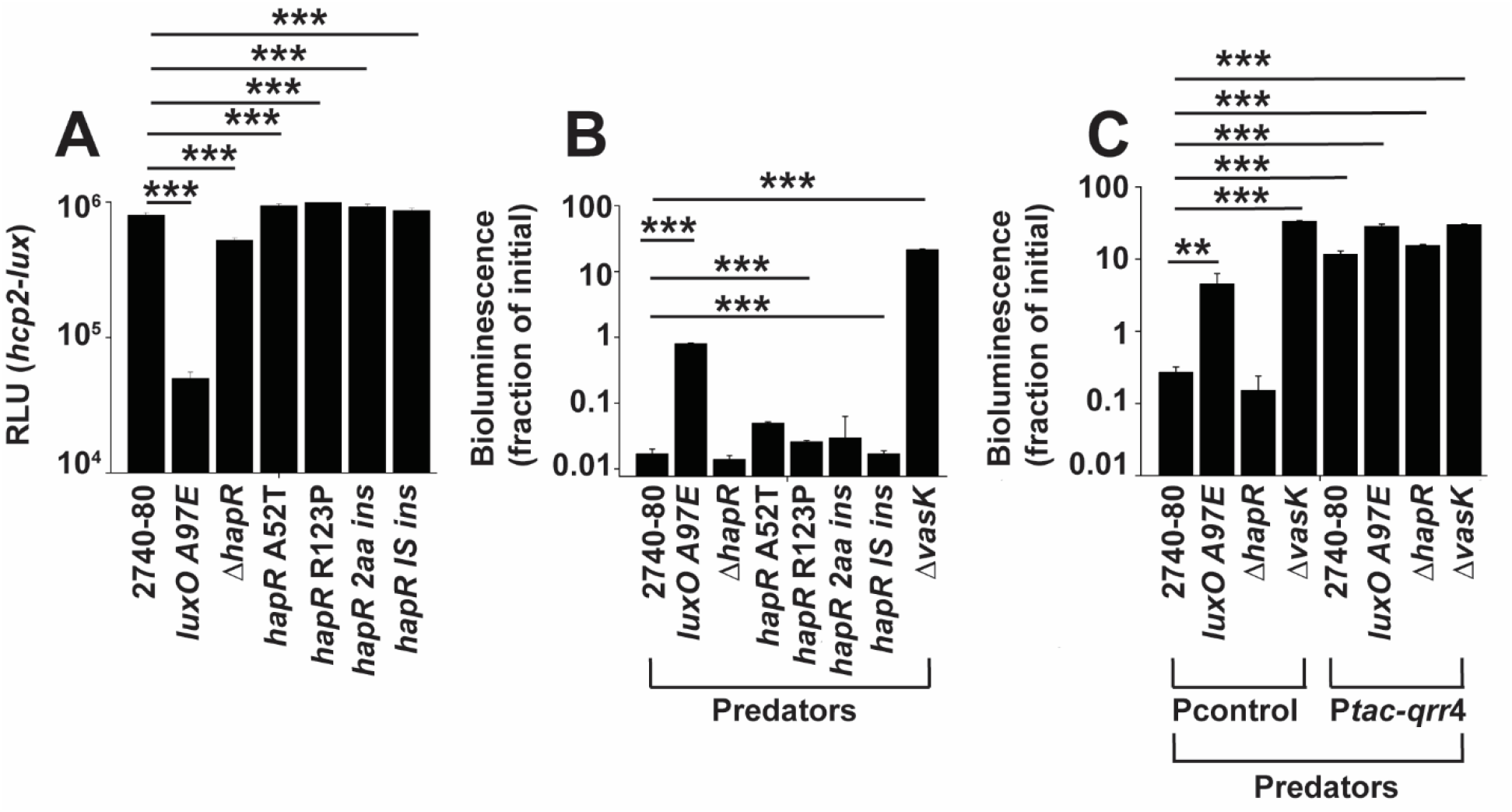
QS control of T6SS-mediated killing activity in *V. cholerae* 2740-80 is driven by LuxO∼P and the Qrr sRNAs independently of HapR. (A) Transcriptional activity of *hcp2*-*lux* in the indicated strains. (B, C) Inter-bacterial T6SS-mediated killing assay measuring survival of *E. coli* Top 10 prey. *E. coli* Top 10 does not possess the T6SS machinery and thus, does not perform T6SS attacks. *E. coli* Top 10 is susceptible to incoming T6SS attacks. *E. coli* Top 10 survival is shown following challenge with the indicated *V. cholerae* predators. The *E. coli* prey cells constitutively express luciferase, so light production can be used as a proxy for live cells (16). In all panels, data represent average values from biological replicates (n = 3), and error bars show SDs. Asterisks indicate statistical significance using a two-tailed Student t test as follows: **, P < 0.005; ***, P < 0.0005. Abbreviations: *hapR* 2aa ins denotes a variant with DNA encoding a two amino acid insertion in *hapR, hapR* IS ins denotes a variant with an IS 200-like element inserted in *hapR*.

To understand whether T6SS-mediated killing activity tracks with level of expression of *t6ss* genes, we measured the capacity of the *V. cholerae* 2740-80 Δ*hapR* strain, the *hapR* variants, and the *luxO A97E* strain to act as predators and kill *Escherichia coli* prey cells in an inter-bacterial T6SS-dependent killing assay. The *E. coli* strain used as the prey in our assay constitutively produces *lux* and is unable to defend itself against incoming T6SS attacks (16). Thus, light output from the *E. coli* prey correlates with live prey cells. To ensure that we are exclusively measuring T6SS-dependent killing, we also assayed a *V. cholerae* 2740-80 strain lacking a T6SS structural protein that is essential for function of the T6SS injection machine, VasK (Δ*vasK*) (23). When the *luxO A97E* strain was used as the predator, there was a ∼50-fold decrease in prey killing relative to when *V. cholerae* 2740-80 was predator (Figure 2B). By contrast, when the Δ*hapR* strain or strains carrying the variant *hapR* alleles were used as predators, they displayed T6SS-mediated killing activity similar to WT *V. cholerae* 2740-80 when it was predator (Figure 2B and Supplementary Table 1). No killing occurred when the Δ*vasK* strain was the predator, confirming that, in our assay, killing requires T6SS activity. Thus, HapR alters neither *t6ss* expression nor T6SS-mediated killing activity in *V. cholerae* 2740-80.

HapR resides at the bottom of the *V. cholerae* QS regulatory cascade, downstream of LuxO and the Qrr sRNAs (Figure 1). Given that LuxO and the Qrr sRNAs are required for T6SS-mediated killing but HapR is not, we wondered how QS control of T6SS-mediated killing occurs in strains lacking HapR or those with attenuated HapR activity. One possibility is that the Qrr sRNAs control T6SS-mediated killing activities by a HapR-independent mechanism. To test this notion, we introduced a plasmid encoding a constitutively expressed representative Qrr sRNA, *qrr*4 (P*tac*-*qrr*4), or the plasmid alone (Pcontrol), into *V. cholerae* 2740-80, the Δ*hapR* strain, and the Δ*vasK* strain and examined their ability to kill *E. coli* in the T6SS-mediated killing assay. Here, we are using the Δ*hapR* strain as the representative for strains with decreased HapR function. Introduction of P*tac*-*qrr*4 into *V. cholerae* 2740-80 and the Δ*hapR* strain resulted in loss of prey killing relative to the strains carrying the control plasmid (Figure 2C). The Δ*vasK* strain displayed no T6SS-mediated killing activity, irrespective of whether it carried P*tac*-*qrr*4 or Pcontrol (Figure 2C). Thus, the Qrr sRNAs repress T6SS-mediated killing function in *V. cholerae* and HapR is dispensable for this activity.

### Despite normal T6SS-mediated killing activity, the *V. cholerae* 2740-80 rim cell death program is abolished in strains lacking HapR

The above results show that HapR does not regulate overall *t6ss* expression nor T6SS activity. Nonetheless, we wondered if HapR plays a role in driving the spatio-temporal pattern of cell death in *V. cholerae* colonies. To explore this possibility, we used a time-lapse fluorescence microscopy assay that we previously developed to track live and dead cell distributions in colonies (16). We assessed colonies of *V. cholerae* 2740-80, *luxO A97E*, Δ*hapR*, and the *hapR* variants. The *luxO A97E* strain lacks the rim cell death program and was included as a control (16). In our assay, live cells are tracked via mKO fluorescent protein produced constitutively from the chromosome of each strain (shown in red). SytoX dye (shown in cyan) marks dead cells. Representative images for *V. cholerae* 2740-80, the *luxO A97E* strain, and the Δ*hapR* strain are shown in Figure 3 to demonstrate how the data are obtained. Ratio-metric data (dead/live cell distributions) are converted into space-time kymographs (Figure 4). In *V. cholerae* 2740-80, the cell death program occurs in two phases. “Phase 1” occurs along the colony rim between ∼8 and 40 h (marked by a white arrow in Figure 3, left panel and a black arrow in Figure 4A). “Phase 2” initiates as a ring in the colony center at ∼44 h, and over the next ∼6 h, cell death propagates inward and outward in an apparent wave (marked by a white arrow in both Figure 3, right panel, and Figure 4A). In contrast to the parent, the *luxO A97E* and Δ*hapR* strains, and each *hapR* variant displayed near absences of Phase 1 cell death along the colony rims (Figure 4; ∼10-fold lower). Each strain exhibited the Phase 2 death pattern at the colony center (Figure 4 and (16)). Thus, despite not altering *t6ss* expression or T6SS-mediated killing function, HapR is required to drive the spatio-temporal cell death pattern at the rims of *V. cholerae* 2740-80 colonies. Because the Δ*hapR* and *hapR* variant colonies phenocopy each other, in the remainder of this work, we focus on the Δ*hapR* strain to understand how HapR influences spatiotemporal cell death.

**Figure 3.**
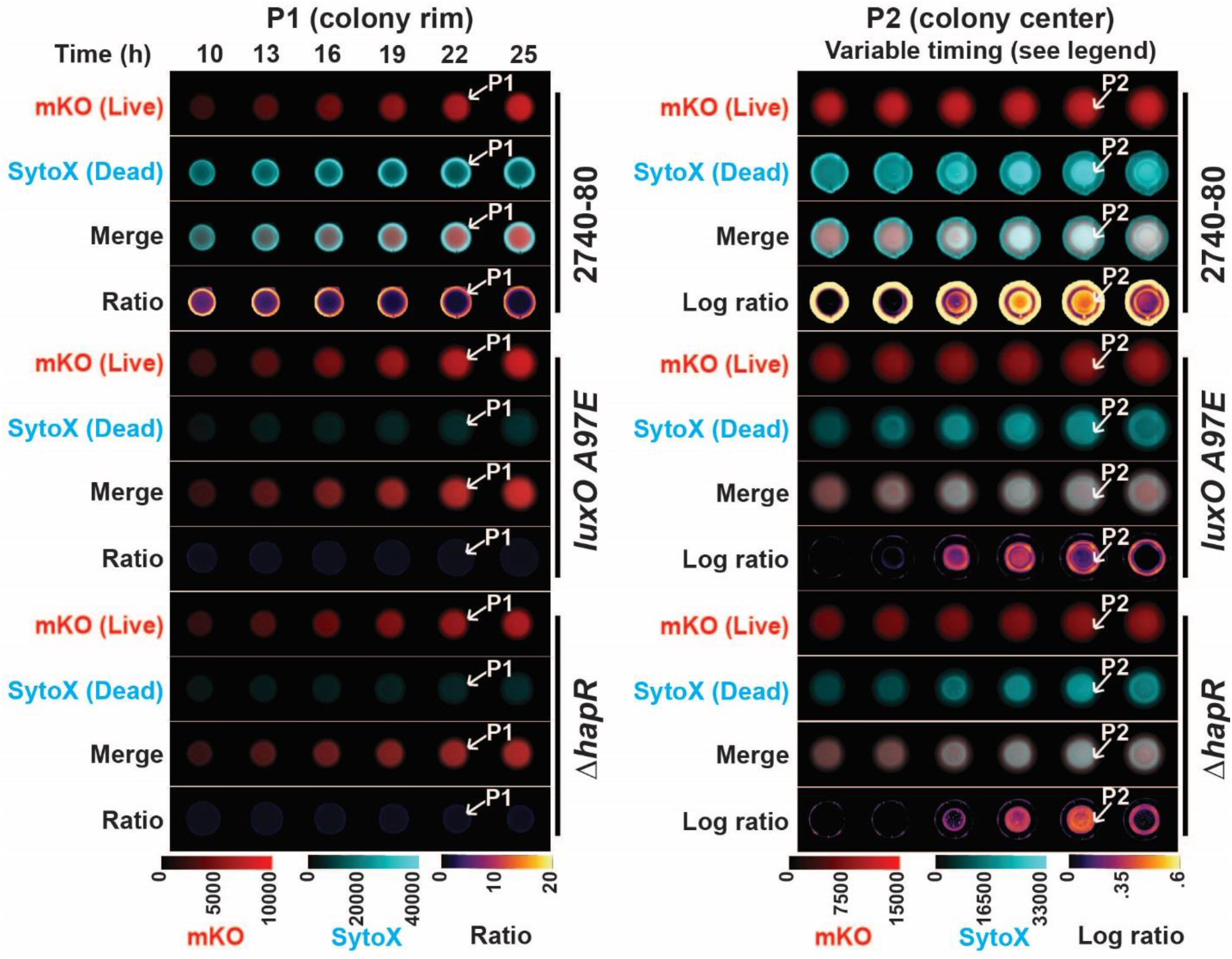
Time-series images demonstrating the two phases of spatio-temporal cell death in *V. cholerae* 2740-80 colonies. Quantitative images from selected time points during growth of the WT *V. cholerae* 2740-80, *luxO A97E*, and Δ*hapR* strains. The strains constitutively produce mKO, which marks live cells. Dead cells are marked with the SytoX stain. The white arrows with the P1 designations highlight the rim of the colony where Phase 1 cell death occurs (left panel). The white arrows with the P2 designations pointing to the colony interior show the Phase 2 cell death ring (right panel). For Phase 2, WT *V. cholerae* 2740-80 colonies are shown between 37-52 h and *luxO A97E* and Δ*hapR* colonies are shown between 27-44 h. There are ∼2.5 h intervals between images, with time increasing from left to right. The differences in timing of Phase 2 among strains has been reported previously (16). Higher level cell death occurs during Phase 1 than Phase 2. Thus, to highlight Phase 2 cell death, logarithmic ratios of the intensities are shown (right panel). For each acquisition channel, the intensity values were mapped using the indicated colors. Scale bars indicate color:intensity.

**Figure 4.**
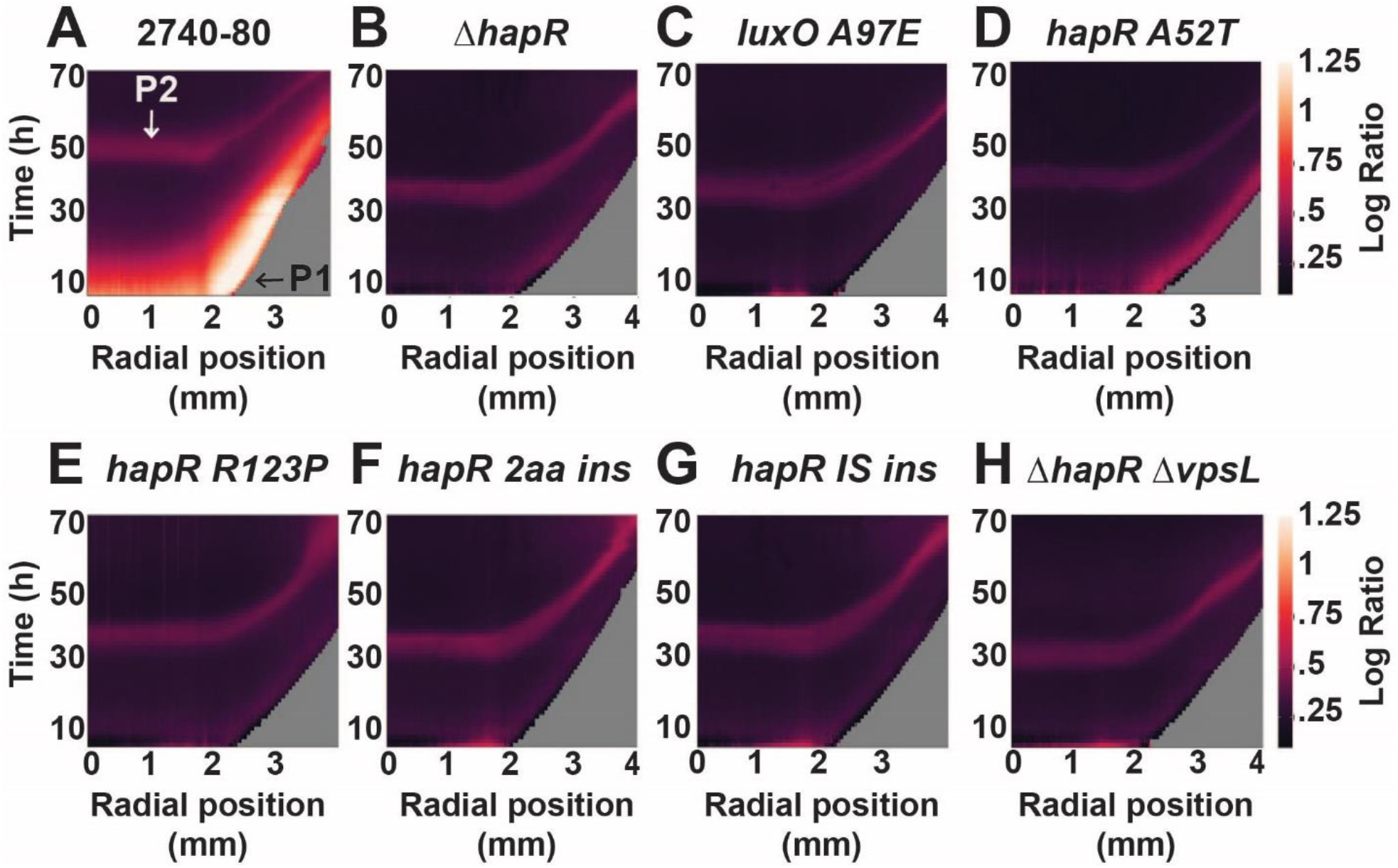
The *V. cholerae* 2740-80 rim cell death program is abolished in a Δ*hapR* strain and in *V. cholerae* 2740-80 *hapR* variants. (A–G) Cell death space-time kymographs of the indicated strains show logarithmic ratio values obtained by dividing the output intensity from the dead-cell channel by that from the corresponding live-cell channel. Ratio values are color-mapped, and the scale bars represent color:intensity. The X axis on each kymograph indicates the radial position in the colony at which the intensity was quantified. The center of the colony is at 0 mm and the colony rim is at ∼3 mm. Phase 1 cell death occurs along the colony rim and is indicated with the black arrow labeled P1 in (A) and cell death is visible as the yellow-colored region. Phase 2 cell death is indicated with the white arrow labeled P2 in (A) and is visible as the red colored region in the colony interior. Kymographs from one colony are presented and are representative of results from ∼3 colonies for each strain.

### Elimination of the ability to form biofilms as a defense against T6SS-mediated killing in the *V. cholerae* 2740-80 Δ*hapR* strain does not restore rim cell death

A mechanism enabling variants in *V. cholerae* 2740-80 colony rims to escape killing is via overproduction of Vps exopolysaccharide that blocks incoming T6SS attacks (16). The Δ*hapR* strain and *hapR* variants exhibit high level *vps* expression. Thus, it is possible that the decreased rim cell death that occurs in the Δ*hapR* strain and *hapR* variants compared to WT *V. cholerae* 2740-80 is a consequence of excess Vps that prevents neighboring cells from engaging in T6SS-mediated killing. If so, we reasoned that a Δ*hapR* strain that is incapable of Vps production would display high colony rim cell death. To test this idea, we tracked cell death in colony rims of a Δ*hapR* Δ*vpsL* strain. VpsL is essential for Vps synthesis. To our surprise, the Δ*hapR* Δ*vpsL* strain had a phenotype identical to the Δ*hapR* strain: minimal death along the colony rim (compare data in Figure 4 panels A and H).

### The *vca0646-0649* operon restores rim cell death in the *V. cholerae* 2740-80 Δ*hapR* strain

In addition to Vps blocking incoming attacks, in *V. cholerae*, T6SS defense is conferred by T6SS immunity proteins, each of which neutralizes one specific T6SS effector toxin protein. Also, a recent Tn-Seq aided genetic screen uncovered several new defense genes that function independently of T6SS immunity proteins, including a gene called *vca0647* (24). The mechanisms by which these components confer T6SS defense remain largely unknown. We wondered whether HapR protects against T6SS-mediated killing at *V. cholerae* 2740-80 colony rims by altering expression of T6SS immunity genes or genes encoding the newly discovered defense proteins. To test this idea, we performed RNA-Seq on WT *V. cholerae* 2740-80 and Δ*hapR* cells isolated from colonies after 20 h of growth, a time when the normal rim cell death pattern is established. Expression of genes encoding T6SS components, including structural, effector, and immunity proteins, was not substantially different in the WT *V. cholerae* 2740-80 and Δ*hapR* strains (see blue in Figure 5). Moreover, most of the recently reported defense gene showed no differences between the two strains (see green in Figure 6). By contrast, the newly identified *vca0647* defense gene displayed higher expression in WT *V. cholerae* 2740-80 than in the Δ*hapR* strain (see red in Figure 5). Our inspection of the DNA sequence surrounding *vca0647* reveals that it resides in a four gene operon (*vca0646, vca0647, vca0648*, and *vca0649*), and indeed, the RNA-Seq data show higher expression of all four genes in WT *V. cholerae* 2740-80 compared to the Δ*hapR* strain (also highlighted in red in Figure 5).

**Figure 5.**
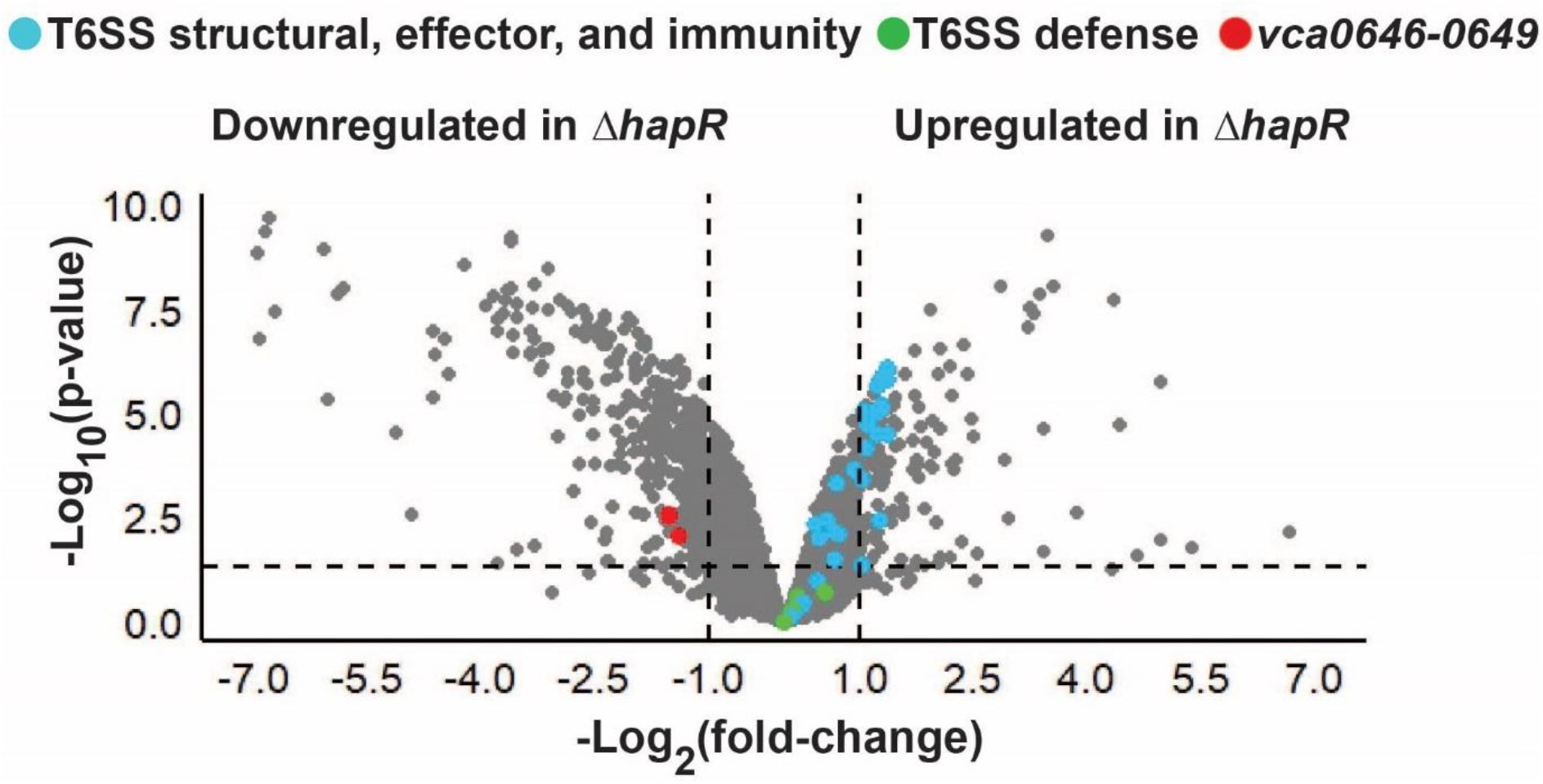
Transcriptomic analysis of *V. cholerae* 2740-80 reveals that HapR controls the *vca0646-0649* operon. Volcano plot displaying fold-changes in gene expression in *cholerae* 2740-80 and Δ*hapR* colonies measured by RNA sequencing. Data are displayed relative to transcript abundance in *V. cholerae* 2740-80. Genes encoding T6SS-mediated killing components (structural, effector, and immunity genes) are highlighted in blue, those encoding T6SS defense components identified by Hersch et. al. (24) are highlighted in green and the *vca0646-0649* genes are highlighted in red. Note: the dots showing the four *vca0646-0649* genes overlap making them difficult to distinguish. Expression levels of individual genes are also provided in Supplementary Table 4. The horizontal line represents a *p*-value of 0.05. Left and right vertical lines represent log_2_ fold-changes of −1 and 1, respectively. Samples are from n*=* 3 biological replicates. Complete datasets are provided in Supplementary Table 4.

**Figure 6.**
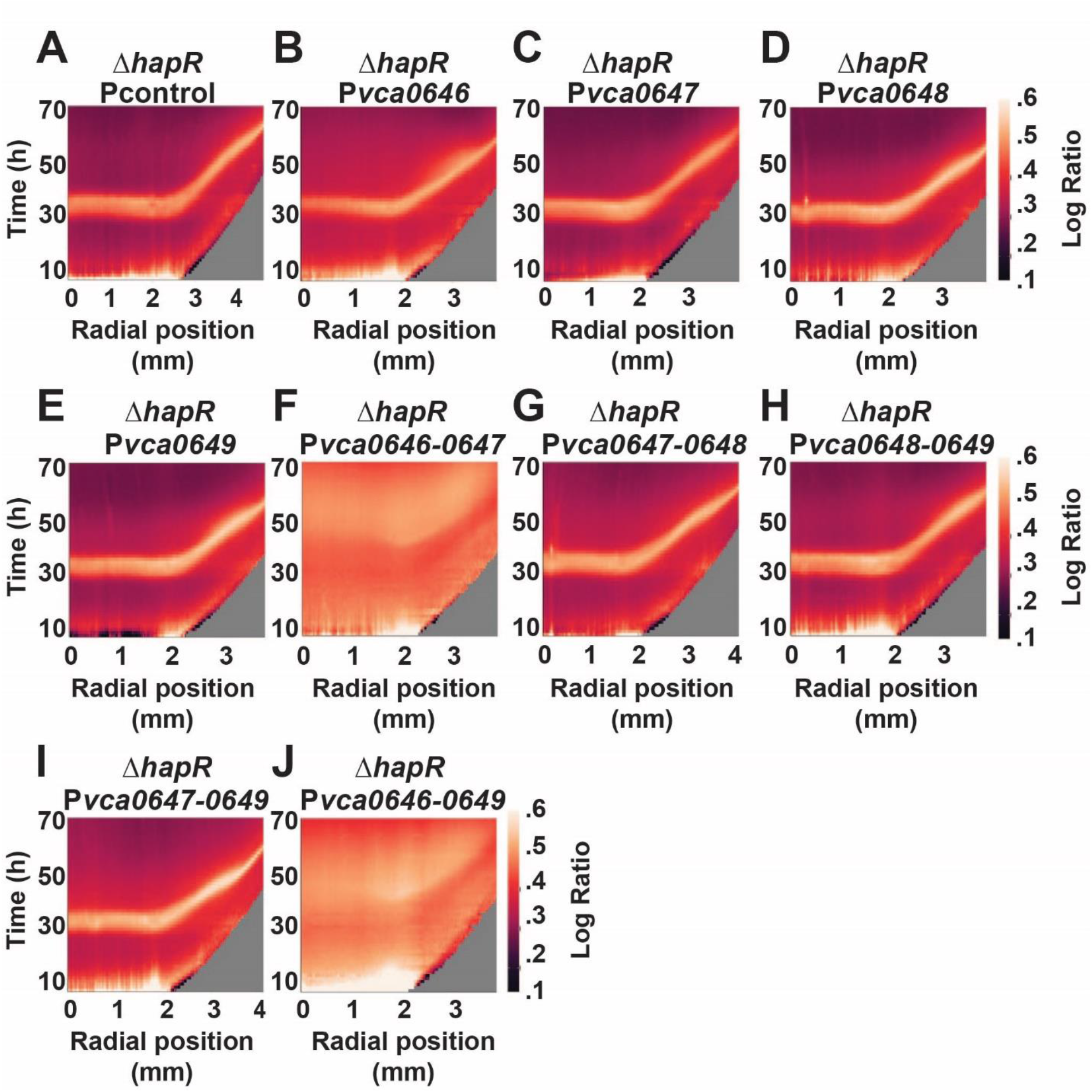
Overexpression of *vca0646-0649* restores rim cell death to the *V. cholerae* 2740-80 Δ*hapR* strain. (A–J) Logarithmic space-time kymographs showing cell death, similar to those in Figure 4, for the indicated strains carrying the designated plasmids. All strains were cultured in the presence of 0.1 % arabinose to induce expression from the P*bad* promoter. Kymographs from one colony are presented and are representative of results from ∼3 colonies for each strain. Companion kymographs for the same strains cultured in the absence of arabinose are provided in Figure S2. Note that the scale used here differs from that in Figure 4. The goal is to enable better visualization of features in strains with low overall cell death.

The *vca0647* gene is predicted to encode a repressor of T6SS defense (24). Based on this earlier report and its high expression in the WT *V. cholerae* 2740-80 strain, we developed the following working model: in the absence of HapR, reduced production of VCA0647 occurs, which enhances T6SS defense and prevents T6SS-mediated killing among cells at the colony rim. If so, we reasoned that increasing the expression of *vca0647* in the Δ*hapR* strain would restore cell death at the colony rim. To test this hypothesis, we introduced a plasmid carrying arabinose-inducible *vca0647* into the Δ*hapR* strain and monitored cell death (P*bad*-*vca0647*). As a control, we introduced an empty vector (Pcontrol). Overexpression of *vca0647* had no effect on the cell death phenotype (Figure 6; compare panels A and C).

The role of *vca0647* in T6SS defense was discovered in a transposon sequencing aided screen, a strategy that can have polar effects on flanking genes. Given that *vca0647* resides in an operon that is more highly expressed in WT *V. cholerae* 2740-80 than the Δ*hapR* strain, we wondered whether the VCA0647 protein acts together with another component(s) encoded in the operon to promote cell death. To explore this possibility, we engineered plasmids carrying different combinations of genes from the *vca0646-0649* operon. Each configuration was placed under control of an arabinose inducible promoter in the Δ*hapR* strain. None of the individual genes modified the Δ*hapR* cell death pattern (Figure 6; compare panels B-E with A). Induction of expression of only one gene pair, *vca0646-0647*, from among three gene pairs tested, increased cell death in the Δ*hapR* strain relative to the control (Figure 6; compare panels F-H with A). Expression of the three gene *vca0647-0649* segment did not change the cell death pattern, whereas expression of the full operon did increase cell death (Figure 6, compare panels I-J with A). Thus, *vca0646* and *vca0647* are both required to influence the cell death program, while *vca0648* and *vca0649* are dispensable.

### *vca0646-0649* activate cell death independently of the T6SS machinery in *V. cholerae* 2740-80

The obvious conclusion from the above findings is that expression of the *vca0646*-*0649* operon restores rim cell death to the Δ*hapR* strain by lowering T6SS defenses. If so, expression of *vca0646*-*0649* in a Δ*hapR* Δ*t6ss* strain, in which cells are incapable of engaging in T6SS mediated killing, would not restore cell death at the colony rim. We engineered a Δ*hapR* Δ*t6ss* strain that lacks all four pairs of T6SS effector-immunity proteins as well as *vasK*, a gene essential for function of the T6SS injection machinery. As expected, cell death did not occur at the rim nor in the center of the Δ*hapR* Δ*t6ss* strain (Figure 7, compare panel C with A). By contrast, introduction of arabinose-inducible *vca0646*-*0649* into the Δ*hapR* Δ*t6ss* strain drove increased overall cell death (Figure 7, compare panel C with panel G and panel D with panel H and see Figure S3 for controls). Indeed, the results resemble those following introduction of the arabinose-inducible *vca0646*-*0649* operon into the Δ*hapR* single mutant that possesses a functional T6SS apparatus (Figure 7, compare panel A with panel E and panel B with panel F). Thus, *vca0646*-*0649* promotes *V. cholerae* cell death by a mechanism that does not rely on the T6SS machine.

**Figure 7.**
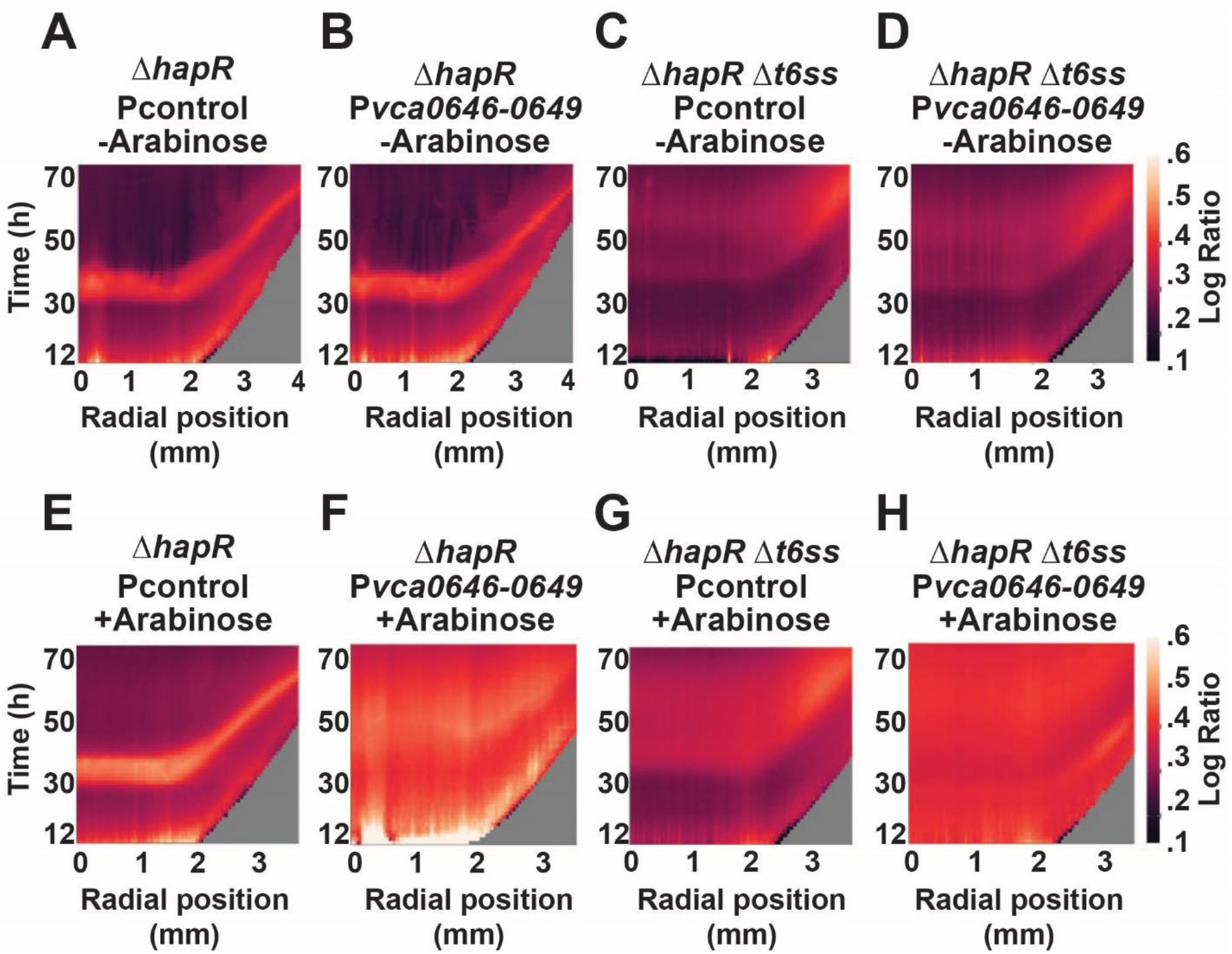
The *vca0646-0649* genes confer cell death to *V. cholerae* 2740-80 independently of the T6SS. (A–H) Logarithmic space-time kymographs showing cell death, similar to those in Figure 4, for the indicated strains carrying the designated plasmids. Strains were cultured in the absence or presence of 0.1 % arabinose to induce expression from the P*bad* promoter. Kymographs from one colony are presented and are representative of results from ∼3 colonies for each strain. Note that the scale used here differs from that in Figure 4. The goal is to enable better visualization of features in strains with low overall cell death. Consequently, it appears as if the Δ*hapR* Δ*t6ss* strain undergoes cell death in the colony center. That is not the case, only residual death occurs relative to that in the WT *V. cholerae* 2740-80 and Δ*hapR* strains as shown in Figure S3.

### Deletion of *vca646-49* does not affect cell death in *V. cholerae*

Given that overexpression of *vca646-49* boosts cell death at the rims of *V. cholerae* colonies (Figures 6 and 7), we reasoned that deletion of *vca0646-0649* would dampen colony rim cell death. We assessed cell death in WT *V. cholerae* 2740-80 and the Δ*vca0646*-*0649*, Δ*t6ss*, and Δ*t6ss* Δ*vca0646*-*0649* strains. Surprisingly, deletion of *vca0646*-*0649* did not alter the cell death patterns (Supplementary Figure 4). Possibly, redundancy exists, and other genes compensate for the loss of *vca0646*-*0649*. Alternatively, *vca0646*-*0649* could function together with some other component and while the high dose of *vca0646*-*0649* delivered by overexpression is sufficient to bypass its requirement, deletion of *vca0646*-*0649* is not.

## Discussion

Here, we discover that QS-regulated spatio-temporal cell death in *V. cholerae* colonies is conferred by at least two pathways operating in parallel (Figure 1). The first cell death pathway, previously described, is driven by the T6SS. The second pathway requires genes in the *vca0646-0649* operon, particularly *vca0646* and *vca0647* but not the T6SS apparatus (Figures 1 and 6). Overexpression of *vca0646-0649* promotes cell death, while deletion of these genes does not influence cell death. This finding suggests redundant or additional components exist that can compensate for loss of *vca0646-0649*. We do not know the functions of any of the VCA0646-0649 proteins. As mentioned, a Tn-Seq study revealed *vca0647* to be a repressor of T6SS defense, but the mechanism was not defined (24). A separate expression analysis reported *vca0646* to be more highly transcribed in classical *V. cholerae* than in El Tor biotypes (25). Our next goal is to discover the functions of VCA0646-0649, with an emphasis on VCA0646 and VCA0647 as well as identify the genes that, in the absence of *vca0646-0649*, drive rim cell death. Another question for future study is whether VCA0646-and VCA0647-mediated cell death is a consequence of self-poisoning or sibling killing.

Our epistasis analyses show that HapR does not regulate *V. cholerae* 2740-80 T6SS-mediated killing (Figure 2). This finding was unexpected because studies conducted in pandemic isolates of *V. cholerae* (C6706 and A1552) demonstrate that HapR activates *t6ss* gene expression (15, 26, 27). A key difference in these T6SS studies may explain our findings. In pandemic isolates, the T6SS machine is not produced under laboratory growth conditions. Environmental stimuli (low temperature or changes in osmolarity) or genetic modification (deletion of *tsrA* encoding a T6SS repressor) are required to induce T6SS-mediated killing in pandemic strains in the laboratory setting (26–28). By contrast, in *V. cholerae* 2740-80, the T6SS system functions during laboratory growth (13, 14). Thus, in the work here, there was no need to expose *V. cholerae* 2740-80 to additional stimuli present in the environment to have T6SS activity. Perhaps, however, in *V. cholerae* 2740-80, HapR only participates in T6SS regulation when the strain is cultured under conditions that closely mimic the environment. Consistent with this logic, the promoter regions driving HapR-controlled T6SS genes (i.e., *vc1415* and *vca0017*) possess 100% sequence identity in pandemic *V. cholerae* C6706 and in *V. cholerae* 2740-80, suggesting that the HapR binding sites are retained in each strain.

The two cell death pathways in *V. cholerae* 2740-80, one that is T6SS-driven and one that is VCA0646-0649-dependent (Figure 1) provide intriguing parallels to cell death mechanisms in higher organisms. In humans, at least five cell death mechanisms exist, each thought to serve a different biological function. For example, apoptosis helps sculpt tissues during development, necroptosis is associated with inflammation and tissue damage, while pyroptosis is relevant during infection or stress (29–32). Perhaps each of the cell death pathways we have discovered in *V. cholerae* is likewise relevant in a specific biological context. For, example, T6SS-mediated cell death could be crucial for development of particular structures such as sectors or biofilm morphological features in bacterial communities. By contrast, the VCA0646-0649 pathway may function in the context of external stress or phage infection by providing a means for members of the community to, respectively, contain the spread of a toxic substance or undergo abortive infection.

## Materials and Methods

### Bacterial growth

*E. coli* S17-1 λ*pir* was used for cloning and conjugations. *V. cholerae* and *E. coli* were cultured in LB medium at 37°C with shaking and with a headspace to growth medium volume ratio of 7. When required, media were supplemented with streptomycin, 200 μg/mL; polymyxin B, 50 μg/mL; kanamycin 50 μg/mL; chloramphenicol, 1 μg/mL. Gene expression was induced with 0.1% arabinose as designated. *V. cholerae* assays were performed at 30°C unless otherwise noted. LB medium, both liquid and solid, was prepared using either dd H_2_O, 100% tap water, or a mixture of 80% tap water and 20% dd H_2_O. These variations in preparation were due to COVID disruptions in supply which made acquisition of LB reagents from multiple vendors necessary. Medium batch differences influenced assay timing and amount of sectoring. However, consistent phenotypes were achieved when solid LB medium was prepared with 80% tap water and 20% dd H_2_O, and liquid LB medium was prepared with 100% tap water (16). Bioluminescence assays were conducted as previously described (16). Relative light units (RLU) denote bioluminescence output divided by culture optical density.

### Strain construction

Chromosomal alterations in *V. cholerae* strains were introduced using either the pKAS32 or pRE112 suicide vectors as previously described (33, 34). Plasmids were constructed using P*bad*-pEVS, pKAS32, or pRE112 as backbones and assembled using the NEB Hi-Fi assembly kit. Plasmids were routinely maintained in *E. coli* S17-1 λ*pir* and introduced into *V. cholerae* strains by conjugation. All strains and plasmids used in the study are listed in Supplementary Tables 2 and 3, respectively.

### Cell death assay

The cell death assay was previously reported (16). Briefly, a 700 μL aliquot of a *V. cholerae* overnight culture was combined with 4 mm glass beads in an Eppendorf tube and the sample subjected to vortex for 5 min to disperse aggregates. The sample was diluted with PBS to reach a final OD_600_ of 0.5. The sample was again subjected to vortex for 5 min, this time without glass beads. A 1 μL aliquot of this suspension was placed onto 35 mL of solid LB agar supplemented with 2 μM SytoX dye (ThermoFisher) in a one well plate and allowed to dry for 5 min at room temperature. The plate was incubated at 30°C for the remainder of the assay. A total of 24 such samples were aliquoted onto each agar pad.

### RNA isolation and sequencing

Strains were cultured on LB agar medium exactly as described for the cell death assay. Subsequently, colonies were resuspended in PBS, 4 mm glass beads were added, and theg suspensions subjected to vortex for 5 min to disperse aggregates. The resulting cell suspensions were treated for 15 min at room temperature with RNAProtect reagent per the manufacturer’s instructions. Thereafter, RNA isolation was performed as described previously (16, 35). Samples were stored at −80°C and shipped on dry ice to SeqCenter (https://www.seqcenter.com/). Sequencing and bioinformatic analyses were conducted as previously described (36). The volcano plot was produced using a custom script in R. Fold-changes for all genes are provided in Supplementary Table 4.

### Image acquisition and analysis

Colonies were plated as described above for the cell death assay. Images of growing colonies were acquired with a Cytation 7 imaging plate reader (Biotek) as reported (16). mKO and SytoX were monitored at ex: 556 and em: 600 nm and ex: 500 and em: 542 nm, respectively. The focal plane was maintained using the Biotek laser autofocus method. At each time point and in each acquisition channel, a 3×3 *xy*-montage of the colony was obtained and stitched together using the linear blend algorithm. A depth of between 225 and 500 μm was sectioned. Maximum intensity *z*-projections were generated for each time point using the Biotek Gen5 software. Fluorescence time-course images of colony growth were analyzed using a custom ImageJ script. First, image background subtraction was performed using a rolling ball radius of 1,000 pixels. Second, to account for shifts during imaging, the sequence of images was registered using the MultiStackReg Fiji plugin and the Rigid Body algorithm. Colony boundaries were determined using the information from the live channel images as a reference and with the aid of the Triangle algorithm. Thereafter, the center of the colony was located with a centroid-finding algorithm using the fluorescent channel that monitored live cells, beginning at the first image acquisition at 8 h and the FeretAngle was determined. The centroid and FeretAngle were used to calculate coordinates to draw a line from the center of the colony to the colony boundary. Finally, spatiotemporal fluorescence intensities in both the live- and dead-cell channels along the line were extracted for kymograph analyses. The regions used to extract intensities were manually monitored to ensure they lacked sectors. The obtained intensity values were used to construct kymograph profiles quantifying the space-time development of live and dead cells within the colony using the R and the ggplot2 visualization packages.

## Data and code availability

Imaging data reported in this study will be shared by the lead contact upon request. Original scripts employed here will be deposited at Zenodo and will become publicly available on the date of publication.

## Acknowledgments

We thank Professor Ned Wingreen for generous feedback about this work and Bassler group members, especially Boyang Qin, for thoughtful discussions. This work was supported by the Howard Hughes Medical Institute, NSF grant MCB-2043238, and NIH grant 5R37GM065859 to B.L.B. A.A.M. is grateful for support from both the Howard Hughes Medical Institute and the Life Sciences Research Foundation through a HHMI-LSRF fellowship.

## Author contributions

A.A.M constructed strains and performed experiments; A.A.M and B.L.B designed experiments and analyzed data; A.A.M wrote custom scripts for image analyses and performed data visualization; A.A.M and B.L.B wrote the manuscript; B.L.B provided oversight, resources, and funding.

## References

1. K. Papenfort, B. L. Bassler, Quorum sensing signal-response systems in Gram-negative bacteria. Nature Reviews. Microbiology 14, 576–588 (2016).

2. C. M. Waters, B. L. Bassler, Quorum sensing: cell-to-cell communication in bacteria. Annual Review of Cell and Developmental Biology 21, 319–346 (2005).

3. M. B. Miller, K. Skorupski, D. H. Lenz, R. K. Taylor, B. L. Bassler, Parallel quorum sensing systems converge to regulate virulence in Vibrio cholerae. Cell 110, 303–314 (2002).

4. Y. Wei, W.-L. Ng, J. Cong, B. L. Bassler, Ligand and antagonist driven regulation of the Vibrio cholerae quorum-sensing receptor CqsS. Molecular Microbiology 83, 1095– 1108 (2012).

5. D. H. Lenz, et al., The small RNA chaperone Hfq and multiple small RNAs control quorum sensing in Vibrio harveyi and Vibrio cholerae. Cell 118, 69–82 (2004).

6. J. D. Mougous, et al., A virulence locus of Pseudomonas aeruginosa encodes a protein secretion apparatus. Science 312, 1526–1530 (2006).

7. R. D. Hood, et al., A Type VI Secretion System of Pseudomonas aeruginosa Targets a Toxin to Bacteria. Cell host & microbe 7, 25–37 (2010).

8. A. B. Russell, et al., Type VI secretion delivers bacteriolytic effectors to target cells. Nature 475, 343–347 (2011).

9. D. L. MacIntyre, S. T. Miyata, M. Kitaoka, S. Pukatzki, The Vibrio cholerae type VI secretion system displays antimicrobial properties. Proceedings of the National Academy of Sciences of the United States of America 107, 19520–19524 (2010).

10. A. B. Russell, S. B. Peterson, J. D. Mougous, Type VI secretion system effectors: poisons with a purpose. Nature Reviews. Microbiology 12, 137–148 (2014).

11. J. Toska, B. T. Ho, J. J. Mekalanos, Exopolysaccharide protects Vibrio cholerae from exogenous attacks by the type 6 secretion system. Proceedings of the National Academy of Sciences of the United States of America 115, 7997–8002 (2018).

12. N. Flaugnatti, et al., Human commensal gut Proteobacteria withstand type VI secretion attacks through immunity protein-independent mechanisms. Nature Communications 12, 5751 (2021).

13. N. C. Drebes Dörr, et al., Single nucleotide polymorphism determines constitutive versus inducible type VI secretion in Vibrio cholerae. ISME J 16, 1868–1872 (2022).

14. S. L. Ng, et al., Evolution of a cis-Acting SNP That Controls Type VI Secretion in Vibrio cholerae. mBio 13, e0042222 (2022).

15. Y. Shao, B. L. Bassler, Quorum regulatory small RNAs repress type VI secretion i n Vibrio cholerae. Molecular microbiology 92, 921–930 (2014).

16. A. A. Mashruwala, B. Qin, B. L. Bassler, Quorum-sensing- and type VI secretion-mediated spatiotemporal cell death drives genetic diversity in Vibrio cholerae. Cell S0092-8674(22)01125–4 (2022). 10.1016/j.cell.2022.09.003.

17. B. Qin, B. L. Bassler, Quorum-sensing control of matrix protein production drives fractal wrinkling and interfacial localization of Vibrio cholerae pellicles. Nat Commun 13, 6063 (2022).

18. M. Simon, M. Silverman, Recombinational Regulation of Gene Expression in Bacteria. (1983). 10.1101/087969176.15.211.

19. G. Koch, et al., Evolution of resistance to a last-resort antibiotic in Staphylococcus aureus via bacterial competition. Cell 158, 1060–1071 (2014).

20. M. Servin-Massieu, Spontaneous appearance of sectored colonies in Staphylococcus aureus cultures. Journal of Bacteriology 82, 316–317 (1961).

21. J. C. van Kessel, L. E. Ulrich, I. B. Zhulin, B. L. Bassler, Analysis of Activator and Repressor Functions Reveals the Requirements for Transcriptional Control by LuxR, the Master Regulator of Quorum Sensing in Vibrio harveyi. mBio (2013). 10.1128/mBio.00378-13.

22. S. T. Miyata, M. Kitaoka, T. M. Brooks, S. B. McAuley, S. Pukatzki, Vibrio cholerae Requires the Type VI Secretion System Virulence Factor VasX To Kill Dictyostelium discoideum ▿. Infect Immun 79, 2941–2949 (2011).

23. S. Pukatzki, et al., Identification of a conserved bacterial protein secretion system in Vibrio cholerae using the Dictyostelium host model system. Proceedings of the National Academy of Sciences of the United States of America 103, 1528–1533 (2006).

24. S. J. Hersch, R. T. Sejuty, K. Manera, T. G. Dong, High throughput identification of genes conferring resistance or sensitivity to toxic effectors delivered by the type VI secretion system. [Preprint] (2021). Available at: 10.1101/2021.10.06.463450v1 [Accessed 20 June 2024].

25. S. Beyhan, A. D. Tischler, A. Camilli, F. H. Yildiz, Differences in Gene Expression between the Classical and El Tor Biotypes of Vibrio cholerae O1. Infect Immun 74, 3633– 3642 (2006).

26. J. Zheng, O. S. Shin, D. E. Cameron, J. J. Mekalanos, Quorum sensing and a global regulator TsrA control expression of type VI secretion and virulence in Vibrio cholerae. Proc Natl Acad Sci U S A 107, 21128–21133 (2010).

27. T. Ishikawa, et al., Pathoadaptive Conditional Regulation of the Type VI Secretion System in Vibrio cholerae O1 Strains. Infect Immun 80, 575–584 (2012).

28. L. Townsley, M. P. Sison Mangus, S. Mehic, F. H. Yildiz, Response of Vibrio cholerae to Low-Temperature Shifts: CspV Regulation of Type VI Secretion, Biofilm Formation, and Association with Zooplankton. Applied and Environmental Microbiology 82, 4441– 4452 (2016).

29. F. K.-M. Chan, N. F. Luz, K. Moriwaki, Programmed necrosis in the cross talk of cell death and inflammation. Annu Rev Immunol 33, 79–106 (2015).

30. Y. Fuchs, H. Steller, Programmed Cell Death in Animal Development and Disease. Cell 147, 742–758 (2011).

31. I. Jorgensen, M. Rayamajhi, E. A. Miao, Programmed cell death as a defence against infection. Nat Rev Immunol 17, 151–164 (2017).

32. J. Yuan, G. Kroemer, Alternative cell death mechanisms in development and beyond. Genes Dev 24, 2592–2602 (2010).

33. M. J. Eickhoff, C. Fei, X. Huang, B. L. Bassler, LuxT controls specific quorum-sensing-regulated behaviors in Vibrionaceae spp. via repression of qrr1, encoding a small regulatory RNA. PLOS Genetics 17, e1009336 (2021).

34. R. A. Edwards, L. H. Keller, D. M. Schifferli, Improved allelic exchange vectors and their use to analyze 987P fimbria gene expression. Gene 207, 149–157 (1998).

35. A. A. Mashruwala, B. L. Bassler, The Vibrio cholerae Quorum-Sensing Protein VqmA Integrates Cell Density, Environmental, and Host-Derived Cues into the Control of Virulence. mBio 11, e01572–20 (2020).

36. A. A. Bridges, J. A. Prentice, C. Fei, N. S. Wingreen, B. L. Bassler, Quantitative input-output dynamics of a c-di-GMP signal transduction cascade in Vibrio cholerae. PLoS Biol 20, e3001585 (2022).

37. L. C. Metzger, et al., Independent Regulation of Type VI Secretion in Vibrio cholerae by TfoX and TfoY. Cell Reports 15, 951–958 (2016).

