## Supplemental Information for "Quorum sensing orchestrates parallel cell death pathways in *Vibrio cholerae* via Type 6 secretion dependent and independent mechanisms"

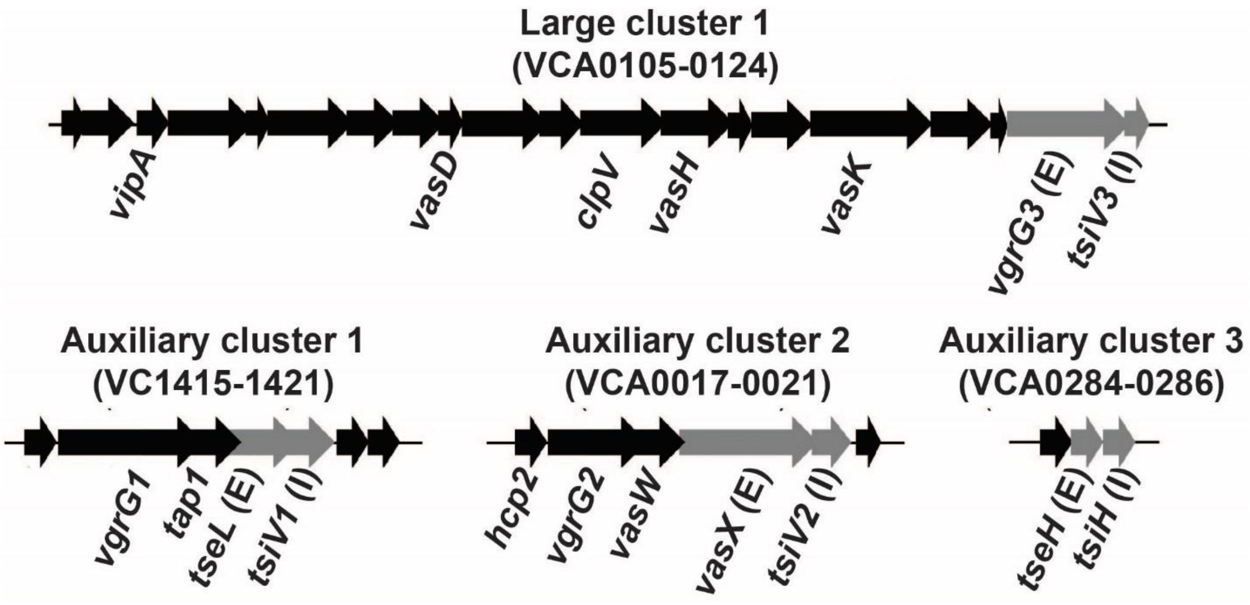


**Supplementary Figure 1. Arrangement of T6SS genes in *V. cholerae* 2740-80.** Select gene names are provided. Genes encoding T6SS effector and immunity proteins are depicted in gray and designated with, respectively, an E or I in parentheses. The large *t6ss* gene cluster is located on the major chromosome and the three auxiliary *t6ss* gene clusters are on the minor chromosome. The large cluster encodes the proteins that make the T6SS secretion complex and one effector-immunity protein pair. Each of the auxiliary clusters encodes one effector-immunity protein pair, among other genes. The figure was adapted from Metzeger et. al. 2016 and Mashruwala and Bassler 2022 (16, 37).

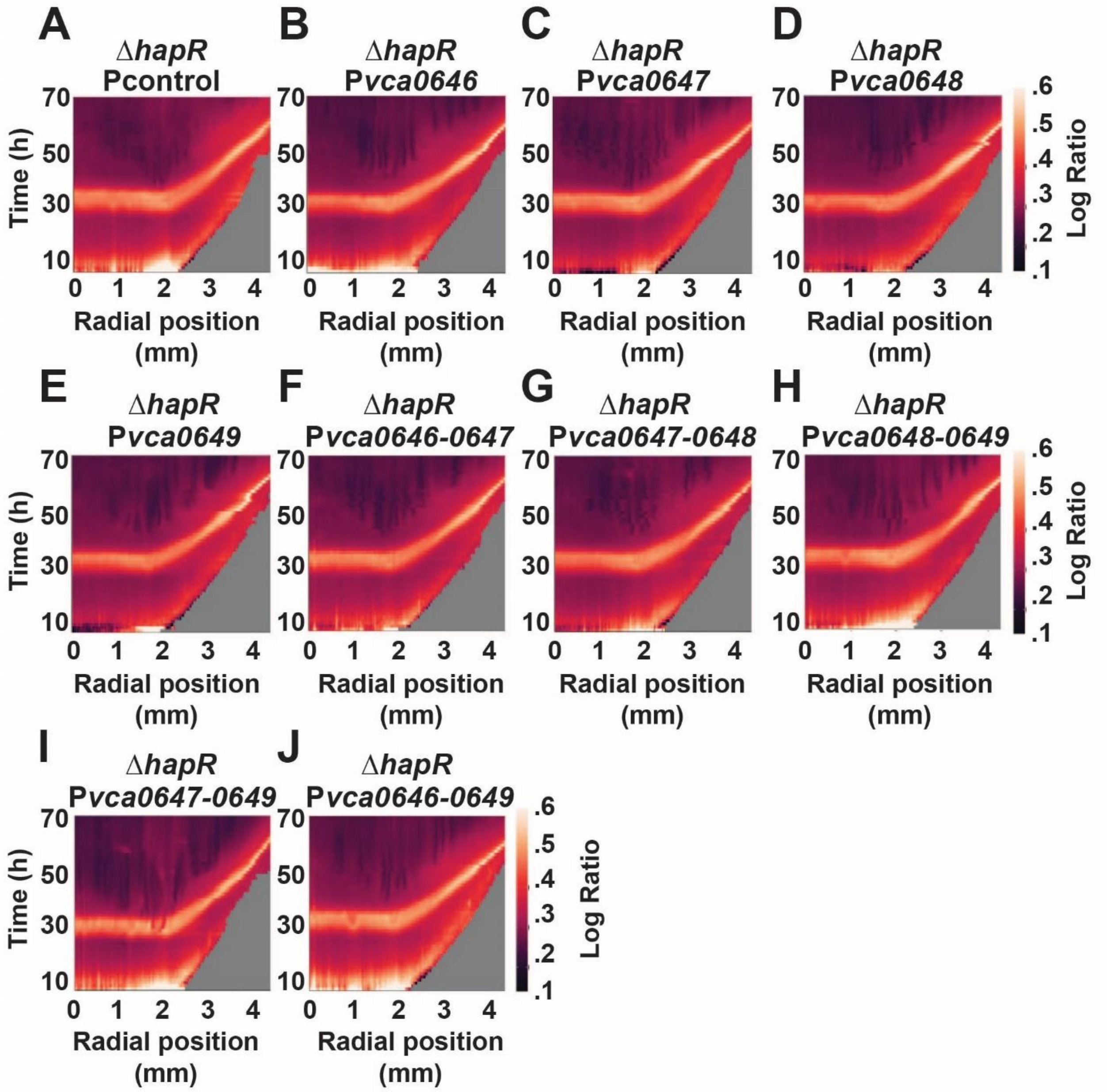


**Supplementary Figure 2: Kymographs showing plasmid controls have no effect on rim cell death in the *V. cholerae* 2740-80 Δ*hapR* strain.** (A–J) Logarithmic space-time kymographs showing cell death for the Δ*hapR* strain carrying the designated plasmids. All strains were cultured in the absence of arabinose. Kymographs from one colony are presented and are representative of results from ~3 colonies for each strain. These data are control kymographs for the same strains presented in Figure 6.

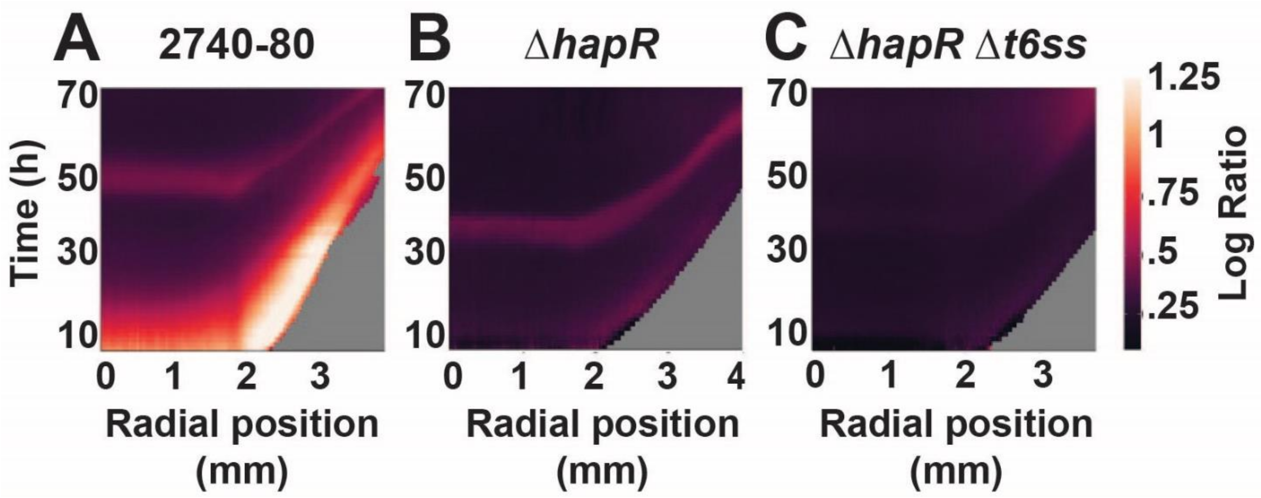


**Supplementary Figure 3: Cell death at the centers of WT *V. cholerae* 2740-80 and Δ*hapR* colonies is driven by T6SS.** (A–C) Logarithmic space-time kymographs showing cell death for the designated strains. Kymographs from one colony are presented and are representative of results from ~3 colonies for each strain. These data accompany Figure 7.

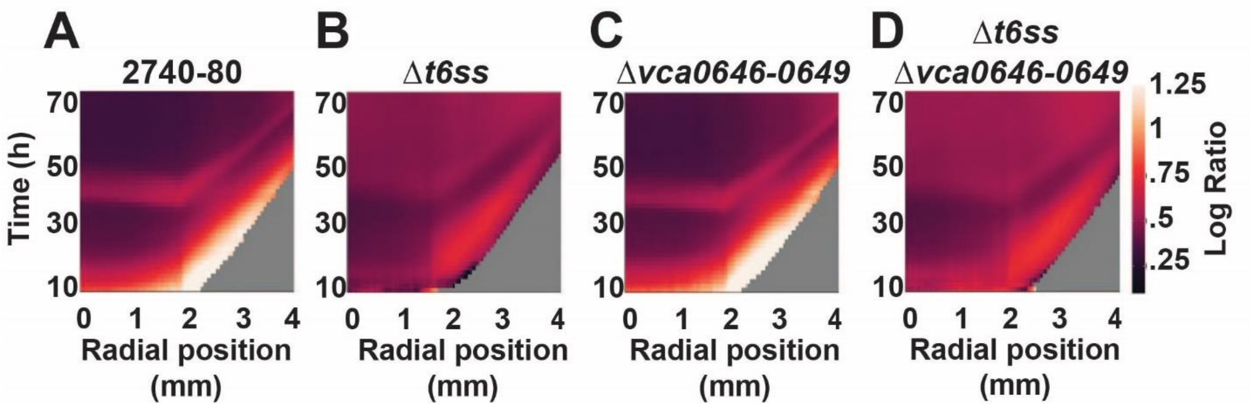


**Supplementary Figure 4: Deletion of *vca646-649* does not alter the *V. cholerae* rim cell death pattern.** (A–D) Logarithmic space-time kymographs showing cell death for the designated strains. Kymographs from one colony are presented and are representative of results from ~3 colonies for each strain.

| **Table S1. Genotypes and phenotypes of QS strains and *hapR* variants used in this study. Data for p*luxC-lux* and p*vpsL-lux* were reported previously and are presented for ease of comparison** (1). | | | | | | |
|---|---|---|---|---|---|---|
| **Genotype** | **Type of mutation** | **p*luxC-lux* transcription: Activated by HapR at HCD. Levels relative to parent.** | **p*vpsL-lux* transcription: Repressed by HapR at HCD. Levels relative to parent.** | **p*hcp2-lux* transcription: Activated by HapR at HCD. Levels relative to parent.** | **T6SS killing: Levels relative to parent.** | **Cell death: Levels relative to parent.** |
| *luxO A97E* | Gain-of-function | Lower | Higher | Lower | Lower | Low rim cell death |
| Δ*hapR* | Loss-of-function | Lower | Higher | Same as parent | Same as parent | Low rim cell death |
| *hapR* A52T | Modification-of-function | Same as parent | Higher | Same as parent | Same as parent | Low rim cell death |
| *hapR* R123P | Attenuation-of-function | Lower | Higher | Same as parent | Same as parent | Low rim cell death |
| *hapR* 2aa insertion (after 54$^{th}$ aa) | Loss-of-function | Lower | Higher | Same as parent | Same as parent | Low rim cell death |
| *hapR* ORF interrupted by IS200/IS605-like element | Loss-of-function | Lower | Higher | Same as parent | Same as parent | Low rim cell death |

| **Table S2. Strains used in this study** | | | |
|---|---|---|---|
| **Strains** | **Genotype/Description** | **Genetic Background** | **Source/Reference** |
| AM-890 | *lacZ*::P*tac-mKO* | 2740-80 | (1) |
| AM-933 | *lacZ*::P*tac-mKO luxO* A97E | 2740-80 | (1) |
| AM-1169 | *lacZ*::P*tac-mKO hapR* A52T; reconstructed variant | 2740-80 | This work |
| AM-1170 | *lacZ*::P*tac-mKO hapR* R120P; reconstructed variant | 2740-80 | This work |
| AM-1171 | *lacZ*::P*tac-mKO hapR* 2aa insertion; reconstructed variant | 2740-80 | This work |
| AM-1172 | *lacZ*::P*tac-mKO hapR* IS200/IS605-like element insertion; reconstructed variant | 2740-80 | This work |
| AM-1130 | *lacZ*::P*tac-mKO* Δ*vasK* | 2740-80 | (1) |
| AM-907 | *lacZ*::P*tac-mKO* Δ*vpsL* | 2740-80 | (1) |
| AM-913 | *lacZ*::P*tac-mKO* Δ*hapR* | 2740-80 | This work |
| AM-1087 | *lacZ*::P*tac-mKO* Δ*hapR* Δ*vpsL* | 2740-80 | This work |
| AM-1229 | *lacZ*::P*tac-mKO* Δ*vca646-0649* | 2740-80 | This work |
| AM-1027 | *lacZ*::P*tac-mKO* Δ*vasK* Δ*vgrG3-tsiV3* Δ*vasX-tsiV2* Δ*tseH-tsiH* Δ*tseL-tsiV1* (Δ*t6ss)* | 2740-80 | (1) |

| | | | |
|---|---|---|---|
| AM-1232 | *lacZ*::P*tac-mKO* Δ*vasK* Δ*vgrG3-tsiV3* Δ*vasX-tsiV2* Δ*tseH-tsiH* Δ*tseL-tsiV1* Δ*vca0646-0649 (*Δ*t6ss* Δ*0646-0649)* | 2740-80 | This work |
| AM421 | *Escherichia coli* | Top10 | Bassler Lab Collection |

| **Table S3. Plasmids used in this study** | | |
|---|---|---|
| **Plasmid name** | **Insert Locus/function** | **Source/Reference** |
| pKAS32 | Suicide vector | Bassler Lab Collection |
| pKAS32-*vpsL* | Construct used to make chromosomal deletion | Bassler Lab Collection |
| pKAS32-*hapR* | Construct used to make chromosomal deletion | Bassler Lab Collection |
| pEVS-pBAD | Cloning vector for arabinose inducible gene expression | Bassler Lab Collection |
| pEVS-pBAD-*vca0646* | Overexpression construct | This work |
| pEVS-pBAD-*vca0647* | Overexpression construct | This work |
| pEVS-pBAD-*vca0648* | Overexpression construct | This work |
| pEVS-pBAD-*vca0649* | Overexpression construct | This work |
| pEVS-pBAD-*vca0646-47* | Overexpression construct | This work |
| pEVS-pBAD-*vca0647-48* | Overexpression construct | This work |
| pEVS-pBAD-*vca0648-49* | Overexpression construct | This work |
| pEVS-pBAD-*vca0647-49* | Overexpression construct | This work |
| pEVS-pBAD-*vca0646-49* | Overexpression construct | This work |
| P*hcp2-luxCDABE* | Luciferase-based *hcp2* transcriptional reporter | This work |
| Ptac-*qrr*4 | Plasmid for overexpression of *qrr*4 under control of the Ptac promoter | Bassler Lab Collection |

### Reference:


1. A. A. Mashruwala, B. Qin, B. L. Bassler, Quorum-sensing- and type VI secretion-mediated spatiotemporal cell death drives genetic diversity in *Vibrio cholerae*. *Cell* S0092-8674(22)01125–4 (2022). https://doi.org/10.1016/j.cell.2022.09.003.
